# Contributions of single-cell mechanics and cell-cell adhesion to multicellular spheroid mechanics

**DOI:** 10.64898/2026.08.04.742605

**Authors:** David Dolgitzer, Eleana Parajón, Douglas N. Robinson, Pablo A. Iglesias

**Author notes:** Correspondence: P.I.; D.D.

## Abstract

Tumor spheroid mechanics arise from both the mechanical properties of individual cells and the adhesive interactions that organize them into tissues. The relative contribution of these two factors to the bulk mechanical behavior, however, remains difficult to disentangle experimentally. Here, we develop a computational model of micropipette aspiration to compare the mechanical response of isolated cells and multicellular spheroids within a common computational framework. By independently varying single-cell stiffness and cell-cell adhesion, we quantify their effects on aspiration dynamics, effective elastic modulus, and viscoelastic relaxation. Our results show that increasing single-cell stiffness substantially alters the mechanics of isolated cells but has limited influence on the effective elastic modulus of multicellular spheroids. In contrast, changes in cell-cell adhesion produce pronounced effects on spheroid effective elastic modulus. Nevertheless, both parameters increase the retardation time governing the transition from the initial elastic response to long-time viscous deformation. These findings suggest that multicellular elasticity is governed primarily by intercellular mechanical coupling, whereas the dynamical response to applied stress depends jointly on cell-scale mechanics and cell-cell adhesion.

## Introduction

Tumor spheroids are multicellular systems whose mechanical properties influence growth, deformation, and invasion [1–3]. During cancer progression, malignant cells form solid aggregates that interact mechanically with their surrounding tissue and may later disseminate through metastatic migration. The mechanical behavior of tumors, including their morphology, response to external stress, and viscoelastic properties, depends both on the mechanical properties of individual cells and on the adhesive interactions that couple neighboring cells together [4–6]. However, the relative contributions of single-cell mechanics and intercellular adhesion to the collective mechanics of tumors remain incompletely understood.

The mechanical response of individual cells is governed by both active and passive processes. Actomyosin contractility, cytoskeletal organization, membrane tension, and intracellular material properties collectively determine a cell’s resistance to deformation [7, 8]. These properties influence migration [9], division [10], interactions with neighboring cells and the extracellular matrix [11, 12], and cellular responses to external stress [10, 13]. Mechanical differences between cells have been associated with distinct cell types, genetic alterations, and stages of cancer progression [14–16]. Because tissues are composed of mechanically active cells, it is natural to expect that single-cell mechanical properties contribute to tissue-scale mechanics. However, cells within a tissue are mechanically coupled through intercellular contacts, making it unclear to what extent tissue-level mechanical behavior can be inferred from measurements performed on isolated cells.

Cell-cell adhesion provides a second major contribution to multicellular mechanics. In epithelial tissues and tumors, adhesion is largely mediated by cadherins, transmembrane proteins that form intercellular junctions through interactions with their those on neighboring cells [17, 18]. These junctions maintain tissue cohesion, transmit mechanical stresses, and regulate the structural organization of multicellular aggregates. Consequently, adhesion influences not only whether cells remain connected, but also how forces are distributed and propagated throughout the tissue. Understanding collective cell mechanics therefore requires distinguishing how individual cells resist deformation from how cells are mechanically coupled to each other.

Computational models have become important tools for studying the mechanics of cells and multicellular tissues, allowing individual mechanical mechanisms to be varied independently under controlled conditions. Vertex models and Cellular Potts Models have been widely used to investigate epithelial mechanics, tissue organization, spheroid growth, and collective cell migration [19–21]. These studies have provided valuable insight into the mechanics of individual cells and the collective behavior of multicellular systems, though often treating these scales separately. Therefore, the relationship between mechanical properties measured at the single-cell level and the mechanical response of multicellular spheroids remains largely unexplored in computational studies. Among the experimental methods used to characterize cellular and multicellular mechanics, micropipette aspiration (MPA) provides a simple and quantitative assay for probing viscoelastic deformation under controlled loading [22, 23]. Because analogous aspiration protocols can be applied to both isolated cells and multicellular spheroids, this technique offers a natural framework for comparing mechanical behavior across organizational scales.

Here, we develop a computational framework to investigate how individual cell mechanics and cell-cell adhesion contribute to the mechanical response of multicellular spheroids. Using the Cellular Potts Model (CPM), we simulate MPA of both isolated cells and spheroids, allowing direct comparison across scales within a common modeling framework. Cellular resistance to deformation is varied through the strength of a perimeter constraint, while intercellular coupling is varied through cell-cell adhesion. We then quantify aspiration dynamics, elastic modulus, and fitted viscoelastic parameters under increasing aspiration forces. Our results indicate that increasing single-cell stiffness strongly affects the mechanical response of isolated cells but has only a limited effect on the elastic resistance of multicellular spheroids. In contrast, cell-cell adhesion substantially influences the elasticity of spheroids. These findings suggest that collective mechanical behavior cannot be inferred directly from single-cell mechanics alone and highlight the importance of intercellular adhesion in determining tissue-scale mechanical properties.

## Methods

### Computational Model

Because our objective was to compare the mechanics of isolated cells with those of multicellular spheroids within a common computational framework, we employed the CPM in both cases. The CPM is a stochastic, lattice-based, and agent-based framework in which cells are represented as extended domains on a discrete lattice [24]. This representation allows individual cellular mechanical properties to be prescribed while simultaneously capturing collective multicellular behavior [25]. To model single-cell and spheroid aspiration, we used Morpheus (version 2.3.9), an open-source modeling and simulation environment for multicellular systems [26]. All simulations were performed on a two-dimensional square lattice using four computational threads. Physical length and time scales were introduced during post-processing rather than during simulation. Lattice units were converted such that a typical cell corresponded to an effective radius of 5 *µ*m, while one Monte Carlo time step was mapped to 0.006 s.

Within the CPM, lattice updates are attempted through stochastic copy operations that evolve according to minimization of an effective Hamiltonian. We used the default “optimal” neighborhood implementation provided by Morpheus. The Hamiltonian is given by

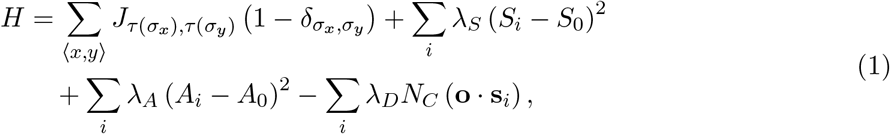

where the first term represents the interfacial contact energy between neighboring lattice sites. Here, *J_σx,σy_* denotes the contact energy between cells identities *σ_x_* and *σ_y_* occupying neighboring sites *x* and *y*, while *τ* (*σ_x_*) and *τ* (*σ_y_*) denote the corresponding cell types. Under this sign convention, lower (more negative) values of *J* correspond to stronger adhesion. This term is zero if only one cell is simulated or if no contact energy is specified. The second term penalizes deviations of the cell perimeter *S_i_* of cell *i* from the target value, 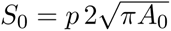, where *p* is a dimensionless scaling factor. We used *p* = 1, corresponding to the perimeter of a circle with area *A*_0_. We refer to *λ_S_* as the shape penalty strength because it controls the effective resistance to changes in cell shape and boundary length. The third term penalizes deviations of the cell area *A_i_* from the target value *A*_0_, with penalty strength fixed at *λ_A_* = 1 throughout the study. Sensitivity analysis indicated that *λ_A_* did not contribute appreciably to the effective elastic modulus over the range considered (Fig. S3). The final term introduces a directed motility bias representing the effective aspiration force. Its magnitude is controlled by *λ_D_*, while *N_C_* denotes the number of lattice sites belonging to cell *i* that lie within the prescribed aspiration region. The vector **s***_i_* describes the displacement associated with an attempted copy event, and **o** specifies the preferred direction of motion.

Copy attempts are accepted according to the Metropolis algorithm,

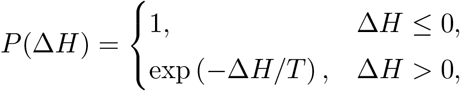

where Δ*H* is the change in Hamiltonian associated with the attempted copy event and *T* is the Monte Carlo sampler temperature controlling the Boltzmann probability of accepting energetically unfavorable updates. Throughout this study, we used *T* = 0.2. Consequently, energetically favorable updates are always accepted, whereas unfavorable updates are accepted with a probability that decreases exponentially with increasing Δ*H*.

The number of attempted copy operations is controlled by the Monte Carlo Step time (*t*_MCS_). We used an *t*_MCS_ duration value of 0.1, corresponding, on average, to 10 update attempts for each lattice site per simulation time step. Boundary conditions were left at the Morpheus defaults, as no explicit boundary effects were required for the simulated geometries.

### Single-cell aspiration

We constructed a cell with a radius of 5 *µ*m (after unit conversion) and simulated MPA by applying an effective aspiration force within the pipette region (Fig. 1a). The pipette walls were represented by two stationary (frozen) cells positioned in parallel to define the desired pipette geometry. These cells had no contact energy with the aspirated cell, serving only as rigid geometric boundaries. Consequently, only the portion of the aspirated cell located inside the pipette experienced the applied effective force, whose magnitude was controlled by *λ_D_*.

**Figure 1:**
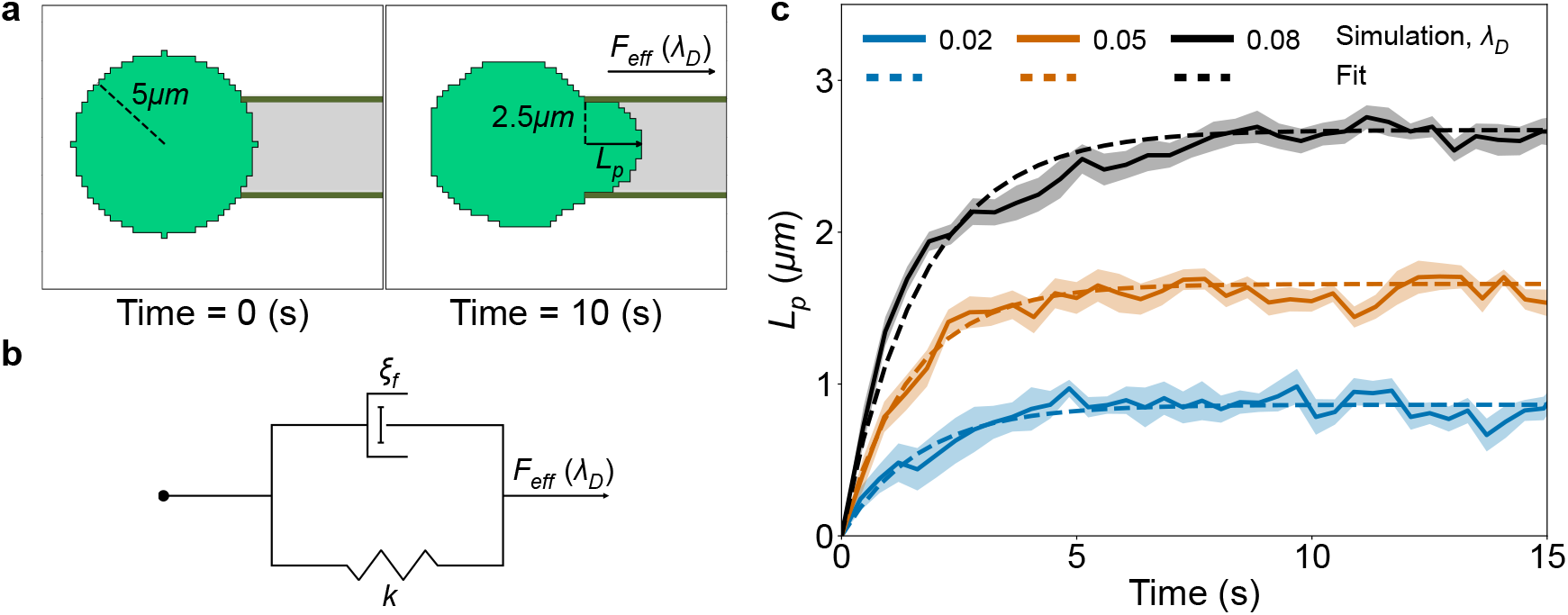
Model and simulation of MPA. **a**. Two-dimensional CPM simulation of single-cell MPA under a constant effective aspiration force parameter *λ_D_*. Each simulation contains a cell with radius 5 *µ*m and a pipette with radius 2.5 *µ*m. The effective force is applied locally within the pipette region (gray). **b**. Kelvin-Voigt model of viscoelasticity used to describe single-cell aspiration dynamics. The spring constant *k* represents the elastic response, and *ξ_f_* represents the effective viscous resistance. **c**. Aspiration length *L_p_* over 10 seconds for three *λ_D_* values. Solid lines and shaded regions represent the mean and standard error of the mean from 10 simulation. Dashed lines represent the model fit according to Eq. (2).

Experimental measurements of cortical tension are typically performed using aspiration pressures close to the Laplace pressure, such that cells undergo an initial deformation followed by resistance to further aspiration [22]. To reproduce this behavior, we considered a range of effective forces for which the simulated cells exhibited finite equilibrium aspiration lengths.

The aspiration dynamics were described using a Kelvin-Voigt viscoelastic model (Fig. 1b),

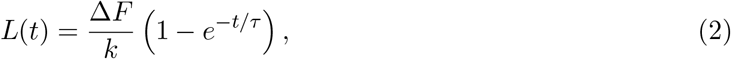

where Δ*F* = *F* − *F*_0_, *F* denotes the total force, *F*_0_ is the minimum force required to produce measurable aspiration, *k* is the effective spring constant, and *τ* = *ξ_f_ /k* is the retardation time. For each effective force, simulations were repeated 10 times and averaged to account for the stochastic nature of the CPM. Increasing the effective force produced progressively larger steady-state aspiration lengths, consistent with the expected Kelvin-Voigt response (Fig. 1c).

### Spheroid aspiration

Following the single-cell aspiration protocol, we constructed spheroids consisting of 500 cells with a chosen cell-cell adhesion strength, controlled by the contact energy parameter *J*. Each spheroid had a radius of 113 *µ*m and was aspirated into a pipette of radius 15 *µ*m (Fig. 3a). Aspiration was simulated by applying an effective force within the pipette region, represented by the localized directed motility term described in Eq. (1). Unlike the single-cell simulations, the effective force was applied dynamically to a one-pixel-thick region at the aspirated front, ensuring that only the leading cells entering the pipette experienced the effective force.

To obtain stable aspiration dynamics under the stochastic CPM framework, several additional modeling assumptions were introduced. First, cells located within a boundary layer of approximately 1.5 cell diameters from the spheroid surface were assigned a reduced shape stiffness (50% of the bulk value). This modification was introduced to reduce the stochastic waiting time before aspiration initiation. Although primarily a computational expedient, this assumption is qualitatively consistent with studies reporting softer peripheral cells in metastatic cancer tumor spheroids [27] and local post-mitotic softening at the spheroid periphery [28]. Second, a temporarily increased effective force was applied until the spheroid had advanced two pixels into the pipette, after which the prescribed effective force was restored. This reduced computational cost by avoiding prolonged stochastic waiting times prior to initial aspiration. Finally, all spheroid cells were assigned a constant contact energy of *J*= −15 with the pipette walls’ interior. Since each wall was represented by a single frozen cell, this interaction maintained contact between the spheroid and the pipette without introducing additional resistance to tangential motion along the wall. The same wall contact energy was used for all spheroid simulations and parametric conditions.

To reproduce the prolonged aspiration observed experimentally in multicellular aggregates, we additionally incorporated Bell-like catch-bond dynamics [29] by allowing cell-cell adhesion to increase exponentially following entry into the pipette, reaching a maximum increase of 50% relative to the baseline adhesion. This local strengthening of adhesion prevented rapid aspiration immediately following entry into the pipette and produced sustained creep consistent with a three-element viscoelastic model consisting of a Kelvin-Voigt element in series with a dashpot (Fig. 3b) [30],

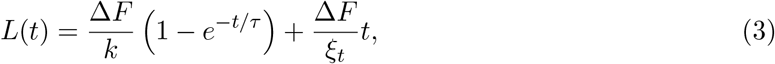

where *ξ_t_*is the long-time viscous coefficient. In the long-time limit,

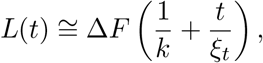

such that the first and second terms represent the elastic and viscous contributions to aspiration, respectively. The aspiration length associated with each simulation was therefore taken as the elastic deformation, *L_p_* = Δ*F*_eff_*/k*, while the second term describes the subsequent viscous creep.

### Cell size normalization

In the single-cell simulations, the aspirated length spans only a few pixels, making the measurements sensitive to lattice discretization. Increasing the simulated cell size reduces this discretization error and improves the precision of aspiration measurements by reducing noise. Conversely, increasing the size of every cell in a 500-cell spheroid substantially increases the computational cost. Consequently, single-cell and spheroid simulations were performed at different lattice resolutions.

To preserve the mechanical behavior across resolutions, we introduced a size normalization of the surface, area, and directed-motion terms in Eq. (1), building on the force characterization of [31]. The resulting Hamiltonian for the single-cell simulations becomes

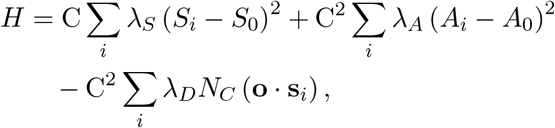

where C = *r*_ref_ */r*_current_, *r*_ref_ denotes the reference cell radius at the chosen lattice resolution, and *r*_current_ is the simulated cell radius.

The *t*_MCS_ time was scaled according to the cell area,

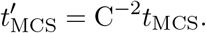

A derivation of the normalization factors and *t*_MCS_ scaling, together with numerical validation demonstrating preservation of the mechanical response across resolutions, is provided in the Supplementary Information.

### Assumptions and computational modifications

The model combines coarse-grained mechanical terms, phenomenological representations of aspiration, and computational procedures introduced to improve simulation tractability. The area, surface, and contact-energy terms constitute the mechanical basis of the CPM and represent effective resistance to deformation and cell-cell coupling. The localized directional bias was used phenomenologically to represent aspiration, because the CPM implementation does not directly include an external force boundary condition.

Because the effective force parameter *λ_D_* was used phenomenologically, the fitted mechanical parameters are reported in CPM units. Accordingly, the fitted spring constant *k* represents effective force units per unit length, whereas *E*_eff_, derived from the slope of effective force versus normalized aspiration length, is expressed in effective force units. These quantities should therefore be interpreted as relative indices of mechanical resistance within the model rather than absolute material moduli.

Additional modifications were introduced specifically for spheroid aspiration. Peripheral cells were assigned a reduced shape penalty strength *λ_S_* to shorten the stochastic waiting time before initial aspiration while preserving the mechanical properties of the spheroid interior. A temporarily increased effective force was applied during the first two pixels of entry for the same computational purpose. In addition, local adhesion strengthening inside the pipette was introduced to prevent immediate unrestricted flow and to reproduce the transition from rapid deformation to long-time creep. This term should be interpreted as a phenomenological representation of force-dependent intercellular stabilization rather than a direct molecular implementation of catch-bond kinetics. All auxiliary modifications were applied identically across the spheroid conditions considered in this study.

### Data Analysis

Simulation outputs were analyzed using MATLAB R2023a, and all figures were generated using Python 3.11.5. Because the simulations are stochastic, trajectories were averaged to calculate the mean and standard error of the mean (SEM), using 10 single-cell trajectories and at least 10 spheroid trajectories for each condition. Trajectories were interpolated onto a common temporal grid using linear interpolation with MATLAB’s interp1 function prior to averaging and model fitting. The models in Eqs. (2) and (3) were then fitted using MATLAB’s lsqcurvefit function, from which the mechanical parameters were extracted.

In a small subset of conditions, mainly in spheroids with softer cells or weaker cell-cell adhesion and under a high effective force value, some spheroid trajectories reached the closed end of the pipette before the end of the 160 s period. To preserve a constant number of trajectories contributed to the average, Eq. (3) was fitted to these trajectories to extrapolate aspiration length to 160 s or, when all trajectories in a condition reached the pipette end prematurely, to the latest endpoint observed within that condition.

At low effective forces, spheroids often remained only partially inserted into the pipette for an extended stochastic waiting period before sustained aspiration began. This initial waiting phase was excluded from the viscoelastic analysis. Instead, aspiration trajectories were analyzed beginning from the onset of sustained aspiration, identified manually from the aspiration length time series as the point after which the spheroid entered continuously into the pipette. The aspiration length and time were then translated such that this point corresponded to *L_p_*= 0 and *t* = 0, respectively. This procedure isolates the viscoelastic response following aspiration initiation and prevents the stochastic entry delay from contributing to the fitted mechanical parameters.

At high effective aspiration force values, some spheroid simulations exhibited a distinct failure mode in which aspiration proceeded too rapidly to maintain a confined protrusion contacting both pipette walls. In these cases, the protrusion became highly deformed and the spheroid translated rapidly toward the end of the pipette. Such trajectories were excluded using a predefined geometric criterion requiring sustained contact with both pipette walls during aspiration.

The threshold effective force, *F*_0_, was estimated from the relationship between the applied effective force and the total aspiration length measured at *t* = 160 s. A linear fit was performed, and *F*_0_ was taken as the force-axis intercept corresponding to zero aspiration. Under the assumed long-time response, aspiration length at any fixed time after the transient relaxation is proportional to *F* − *F*_0_. Therefore, the theoretical intercept is independent of the selected late-time measurement point, although its numerical estimate may vary because of stochastic variability and deviations from the fitted model.

### Experimental Spheroid Micropipette Aspiration

Panc10.05 cells were used to generate spheroids. MPA was performed with an instrumental setup as previously described in [32]. In short, micropipettes of ∼50 *µ*m radius were forged using the MDI Programmable Multipipette Puller (SKU BZ10133958). The exact radius of each pipette was measured for each experiment. Spheroids were collected with a P1000 in order to not disrupt them and placed in cell culture media on imaging chambers. A small aspiration/negative pressure was applied on the system (around 10 mm height on the water level tank or 0.098 nM*/µ*m^2^). This is the starting pressure that allows the spheroid to form attachment between the spheroid and the pipette tip. Each starting pressure will be slightly different for each spheroid. At frame 1 of collection, the acquisition was paused, and then the water level tank height was changed to 100 mm or the pressure was changed to 0.098 nM*/µ*m^2^ while image acquisition was resumed (it took ∼ 35 s to reach the final pressure). Time lapse images (5 s/frame) capturing spheroid deformation were acquired on an Olympus IX81 microscope equipped with MetaMorph software. Microscopy images were analyzed in Fiji (ImageJ). Aspiration was tracked using a straight line drawn parallel to the pipette axis, extending from the pipette entrance to the tip of the aspirated protrusion. The aspiration at frame 1 was subtracted from all subsequent measurements.

## Results

### Shape constraint correlates to cellular elasticity

The cellular shape penalty strength *λ_S_* was evaluated as a potential effective stiffness parameter by comparing three values, *λ_S_* = 0.8, 1.0, and 1.2. Aspiration dynamics were analyzed over a range of six forces, selected such that cells reached a finite equilibrium aspiration length consistent with the Kelvin-Voigt model (Fig. 1b, Eq. (2)). As shown in Fig. 2a, all three cell types exhibit comparable aspiration at low forces. As the applied force increases, however, stiffer cells consistently aspirate less into the pipette. This trend is also evident visually (Fig. 2b, *λ_D_* = 0.08), where cells with lower *λ_S_* protrude further into the pipette. Together, these observations indicate that increasing *λ_S_* increases the effective resistance of the cell to deformation, supporting its interpretation as an effective stiffness parameter within the CPM.

**Figure 2:**
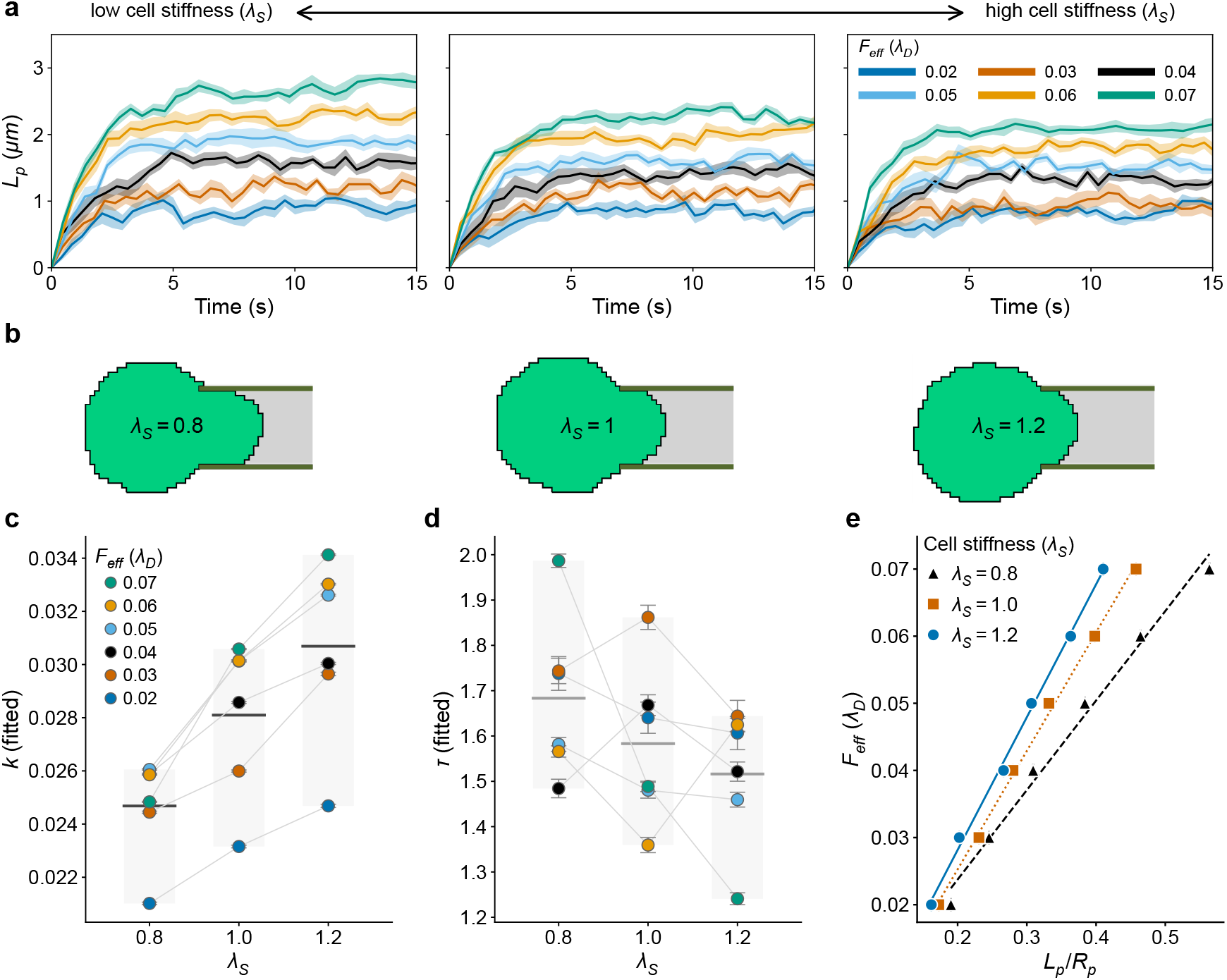
Single-cell mechanics controlled by the shape penalty strength. *λ_S_***. a**. Single-cell aspiration length *L_p_*over time for six values of the effective aspiration force parameter *λ_D_* across three values of shape penalty strength *λ_S_*: 0.8 (left), 1.0 (middle), and 1.2 (right). **b**. Simulation snapshots of three aspirated cells corresponding to the three *λ_S_*values under the same force (*λ_D_* = 0.08). **c**. Fitted spring parameter *k* for the three *λ_S_* values. Gray lines connect values obtained at the same force. **d**. Fitted retardation time *τ* for the three *λ_S_* values. Gray lines connect values obtained at the same force. **e**. Aspiration force as a function of normalized aspiration length *L_p_/R_p_* for the three *λ_S_* values. Linear regression slopes are proportional to the effective elastic modulus, 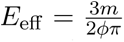. Effective elastic moduli are 0.03 (*λ_S_* = 0.8), 0.04 (*λ_S_* = 1.0), and 0.045 (*λ_S_* = 1.2). Error bars indicate 95% confidence intervals estimated from nonlinear least-squares regression.

**Figure 3:**
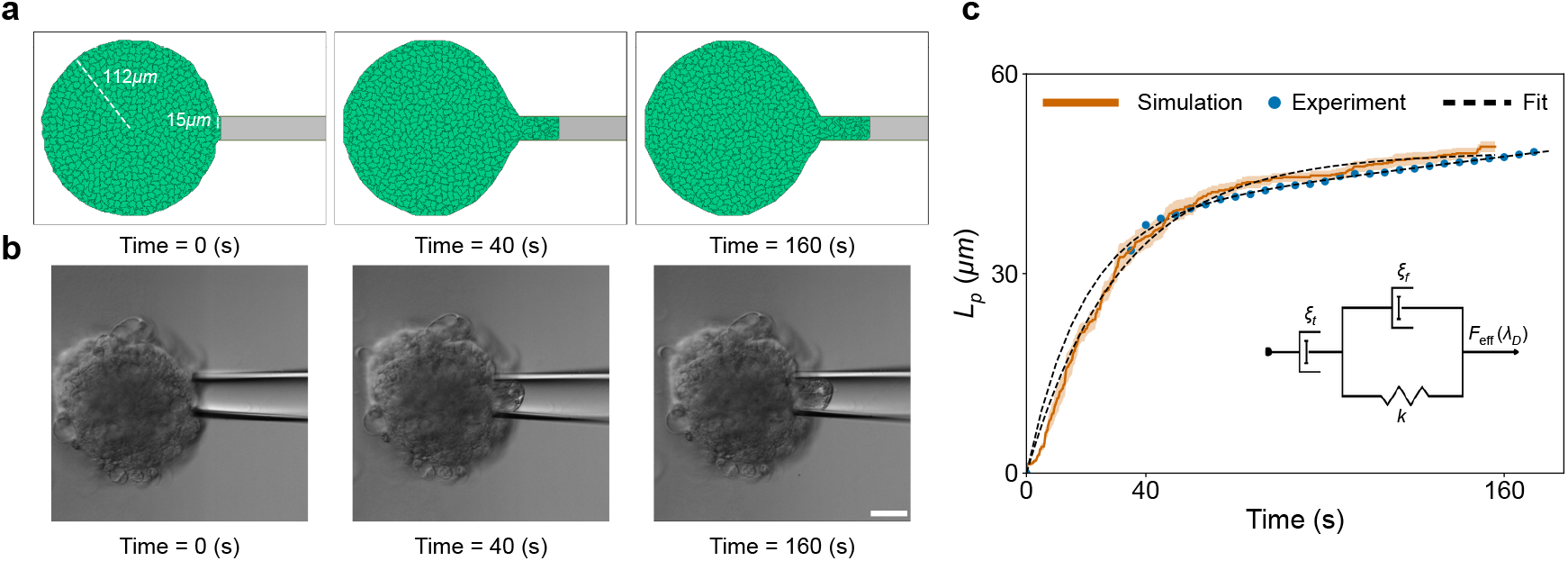
Representative spheroid aspiration in experiment and simulation. **a**. Simulated micropipette aspiration of a multicellular spheroid before aspiration (left), after 40 s (middle), and after 160 s (right). The simulated spheroid had a shape penalty strength of *λ_S_* = 1.2 and cell-cell contact energy *J* = 60, with a force of *λ_D_* = 0.7. **b**. Corresponding experimental images of a multicellular spheroid before aspiration and at the same time points. Scale bar: 50 *µ*m. **c**. Aspiration length as a function of time for the representative simulation and experiment. The solid line and shaded region show the mean and standard error of 10 simulation trajectories, while blue points show the experimentally measured aspiration length. Dashed lines show fits of the three-element viscoelastic model in Eq. (3).

To characterize the aspiration dynamics parametrically, we fitted Eq. (2) to each trajectory and extracted the spring constant *k* (Fig. 2c). Within each stiffness condition, *k* increased for the first few forces before approaching an approximately constant value. More importantly, increasing *λ_S_* consistently resulted in larger values of *k* across the entire range of forces, indicating increased resistance to deformation. Cells with *λ_S_* = 0.8 exhibited a mean *k* of 0.025, increasing to 0.028 for *λ_S_*= 1.0, and 0.031 for *λ_S_* = 1.2.

The same model fits were used to estimate the retardation time *τ* (Fig. 2g). In contrast to the trend observed for *k*, the mean *τ* decreased slightly with increasing *λ_S_*, potentially indicating faster relaxation in stiffer cells. However, the values obtained at individual aspiration forces showed substantial stochastic variability and did not follow this trend consistently. Cells with *λ_S_* = 0.8 exhibited a mean *τ* of 1.68 s, decreasing to 1.58 s for *λ_S_* = 1.0, and 1.52 s for *λ_S_* = 1.2.

Following the classical MPA formulation [22], we plotted the aspiration force against the normalized aspiration length, *L_p_/R_p_*, to determine the effective elastic modulus (Fig. 2e). In all three cases, the relationship is approximately linear, and linear regression was used to determine the corresponding slopes. These were converted to elastic moduli according to 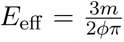, where *ϕ* ≈ 2.1 is an empirical correction factor [22]. Cells with *λ_S_*= 0.8 exhibited the lowest elastic modulus (0.03), followed by *λ_S_* = 1.0 (0.04) and *λ_S_* = 1.2 (0.045). This increase in elastic modulus with increasing *λ_S_* demonstrates that the latter provides a controllable measure of single-cell stiffness within the model.

### Experimental benchmarking of simulated spheroid aspiration

Before examining the effects of cellular stiffness and cell-cell adhesion on spheroids, we compared the simulated aspiration response with a representative experimental MPA trajectory from a Panc10.05 spheroid. Both the experimental spheroid and the simulated spheroid exhibited an initially rapid increase in aspiration length followed by slower, sustained creep over the remainder of the observation period, characteristically to a viscoelastic material under load (Fig. 3a,b). In addition, the simulation reproduced the approximate aspiration magnitude and characteristic time course of the experimental trajectory (Fig. 3c), and both curves were fitted using the same three-element viscoelastic model in Eq. (3) (diagram in Fig. 3c). This comparison was used as a benchmark confirming that our model generated an experimentally-consistent spheroid aspiration regime.

### Cell stiffness has a limited affect on spheroid elasticity

Having established that increasing the shape penalty strength *λ_S_*increases the effective elastic modulus of isolated cells, we next examined how this mechanical property propagates to multicellular spheroids. Three spheroid types were constructed, each consisting exclusively of one of the three cellular stiffness conditions considered previously. The resulting aspiration dynamics consisted of a rapid initial deformation, lasting approximately 40 s (Fig. 3c, middle panel), followed by a slower linear creep into the pipette, a behavior consistent with the three-element viscoelastic model in Eq. (3).

The simulated aspiration dynamics are shown in Fig. 4a for spheroids composed of cells with increasing single-cell stiffness (left to right). Five forces were applied to each spheroid type. Increasing the cellular stiffness reduced both the initial elastic deformation and the total aspiration after 160 s. These differences also corresponded to distinct threshold forces, *F*_0_, obtained from the total aspiration lengths. Spheroids composed of *λ_S_*= 0.8 cells exhibited *F*_0_ = 0.05, increasing to *F*_0_ = 0.17 for *λ_S_* = 1.0 cells and *F*_0_ = 0.20 for *λ_S_* = 1.2 cells. In addition, the transition from the initial elastic response to the long-time viscous regime became progressively longer as cellular stiffness increased. This was seen more clearly for lower force values.

**Figure 4:**
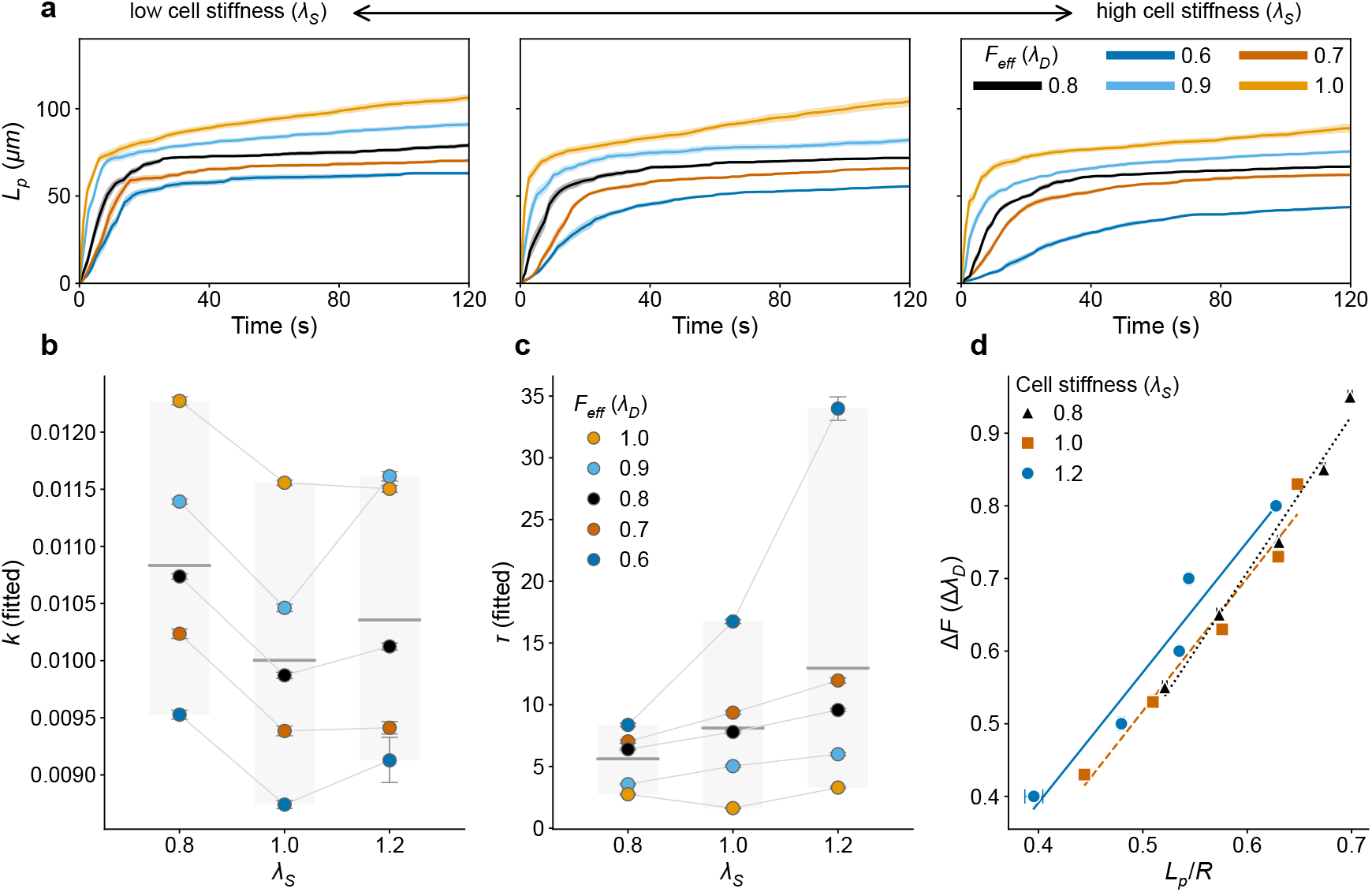
Spheroid aspiration mechanics with varying single-cell stiffness. **a**. Spheroid aspiration length *L_p_* over time for five values of effective aspiration force parameter *λ_D_* across three *λ_S_* values: 0.8 (left), 1.0 (middle), and 1.2 (right). **b**. Fitted spring parameter *k* values for three *λ_S_*values. Gray lines connect values with the same force. **c**. Fitted retardation time *τ* values for the three *λ_S_* values. Gray lines connect values with the same force. **d**. Aspiration force as a function of aspiration length normalized by the spheroid radius for three *λ_S_* values. The slope of the linear fits is proportional to the effective elastic modulus, 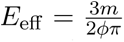 . Effective elastic moduli are 0.49 (*λ_S_* = 0.8), 0.42 (*λ_S_* = 1.0), and 0.42 (*λ_S_* = 1.2). Error bars indicate 95% confidence intervals estimated from nonlinear least-squares regression.

The fitted spring constant *k* similarly showed only minor variation with cellular stiffness (Fig. 4b). Spheroids composed of *λ_S_* = 0.8 cells exhibited a mean *k* of 0.011, compared with approximately 0.010 for spheroids composed of *λ_S_* = 1.0 and *λ_S_*= 1.2 cells. Within each spheroid type, however, *k* increased with increasing force, more consistently than the trend observed for isolated cells.

The fitted retardation time *τ* increased with cellular stiffness despite the opposite trend observed for isolated cells (Fig. 4c). Within each adhesion condition, however, increasing the force reduced *τ* . The mean value of *τ* increased from 5.6 s for spheroids composed of *λ_S_* = 0.8 cells to 8.1 s and 13.0 s for spheroids composed of *λ_S_* = 1.0 and *λ_S_* = 1.2 cells, respectively. Within each spheroid type, increasing the force reduced *τ* . Thus, although stiffer individual cells relaxed more rapidly in isolation, spheroids composed of those same cells exhibited slower collective relaxation.

Finally, we determined the effective elastic modulus by plotting the applied force against the normalized aspiration length, *L_p_/R*, where *R* is the spheroid radius (Fig. 4e). The relationship remained approximately linear for all three spheroid types, allowing the elastic modulus to be obtained using the same procedure employed for isolated cells. Despite the clear increase in elasticity observed at the single-cell level, the bulk elastic modulus of the spheroids remained nearly unchanged. Spheroids composed of *λ_S_* = 0.8 cells exhibited an elastic modulus of 0.49, compared with 0.42 for spheroids composed of both *λ_S_*= 1.0 and *λ_S_*= 1.2 cells. These results indicate that increasing the mechanical stiffness of individual cells alone had little influence on the elastic modulus of the spheroid.

### Cell-cell adhesion enhances spheroid elasticity

Cell-cell adhesion is a key determinant of multicellular organization and mechanical integrity. Having found that varying cellular stiffness produced only minor changes in the elastic modulus of spheroids, we next examined whether altering the mechanical coupling between neighboring cells has a greater influence on the bulk response. To this end, we considered spheroids composed of identical cells (*λ_S_* = 1) under three adhesion strengths, *J* = −40, −50, and −60, where increasingly negative values correspond to stronger cell-cell adhesion.

The aspiration dynamics for the three adhesion conditions are shown in Fig. 5a. The same five forces were applied as in the previous section. In contrast to varying the cellular stiffness, changes in cell-cell adhesion produced substantially different aspiration profiles. Increasing adhesion reduced both the initial elastic deformation and the total aspiration length, while also decreasing the sensitivity of aspiration to increasing force values. Consequently, strongly adhesive spheroids exhibited comparatively small changes in aspiration over the investigated force range, whereas weakly adhesive spheroids showed progressively larger aspiration with increasing force. These differences were reflected in the corresponding estimated threshold force *F*_0_, obtained from the total aspiration lengths at 160 s. Spheroids with *J* = −40 exhibited *F*_0_ = 0.26, decreasing to *F*_0_ = 0.17 for *J* = −50, while no measurable threshold was observed for *J* = −60 (*F*_0_ = 0). As in the previous section, increasing adhesion also prolonged the transition from the initial elastic deformation to the long-time viscous regime.

**Figure 5:**
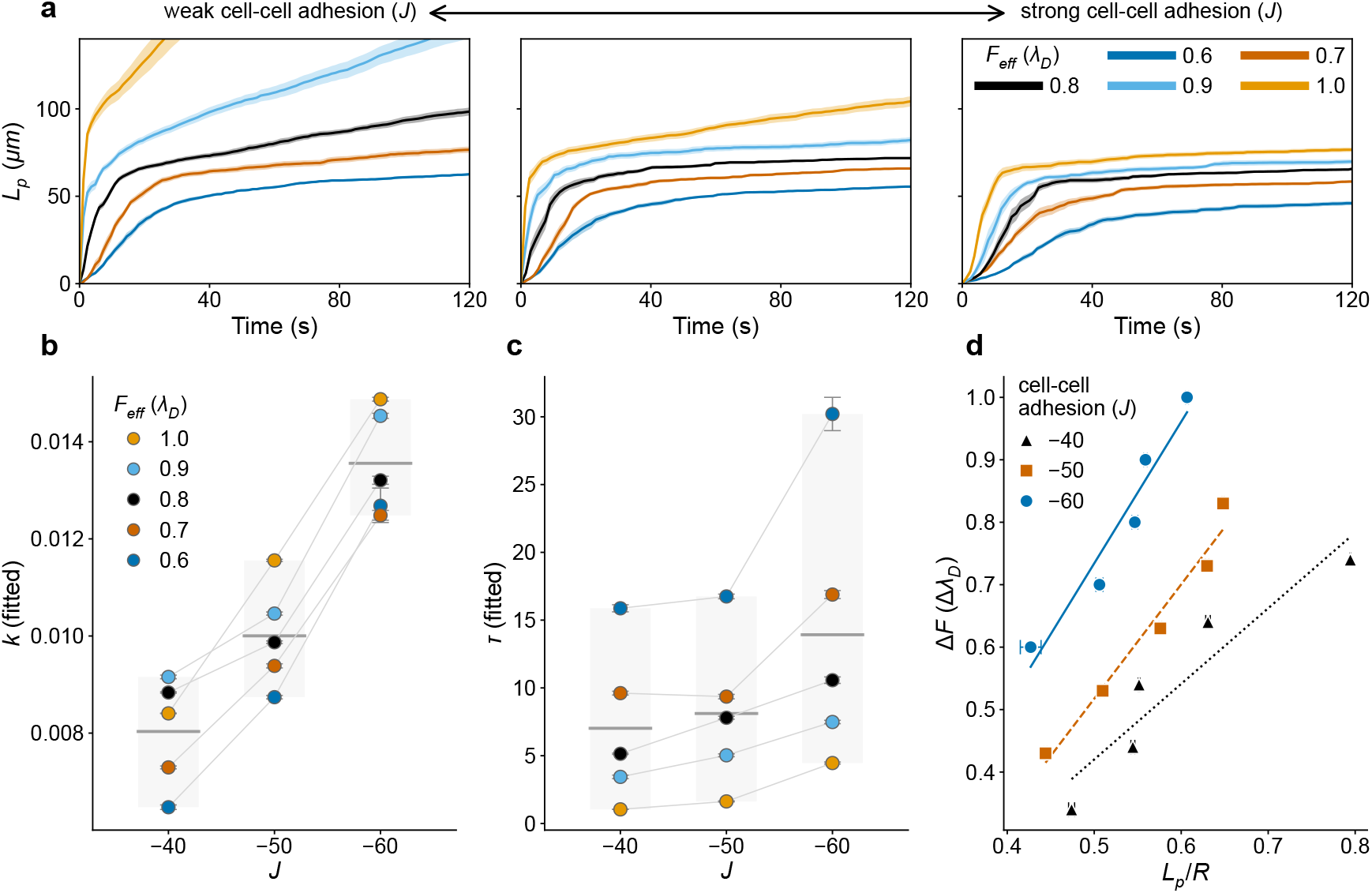
Spheroid aspiration mechanics with varying cell-cell adhesion. **a**. Spheroid aspiration length *L_p_* over time for five values of effective aspiration force parameter *λ_D_* across three adhesion *J* values: 40 (left), 50 (middle), and 60 (right). **b**. Fitted spring parameter *k* values for three *J* values. Gray lines connect values with the same force. **c**. Fitted retardation time *τ* values for three *J* values. Gray lines connect values with the same force. **d**. Aspiration force as a function of aspiration length normalized by the spheroid radius for three *J*values. The slope of the linear fits is proportional to the effective elastic modulus, 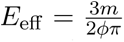. Effective elastic moduli are 0.28 (*J* = 40), 0.42 (*J* = 50), and 0.52 (*J* = 60). Error bars indicate 95% confidence intervals estimated from nonlinear least-squares regression.

We next fitted Eq. (3) to the aspiration trajectories and extracted their spring constant *k* (Fig. 5b). Although *k* again increased with increasing force within each adhesion condition, varying the adhesion strength produced a much stronger effect than varying the cellular stiffness. The mean value of *k* increased from 0.008 for spheroids with *J* = −40 to 0.010 for *J* = −50, reaching 0.014 for *J* = −60, indicating that stronger mechanical coupling between neighboring cells increases the stiffness of the aggregate.

The fitted retardation time *τ* was also evaluated to examine how adhesion influences the collective relaxation dynamics (Fig. 5c). Similarly to spheroids comprised of cells with different stiffness, *τ* reduced with increasing force within each adhesion condition. Across the three spheroid types, however, stronger adhesion consistently increased the mean retardation time, from 7.0 s for *J* = −40 to 8.1 s and 13.9 s for *J* = −50 and *J* = −60, respectively.

Finally, the effective elastic modulus was determined from the relationship between force and normalized aspiration length, *L_p_/R* (Fig. 5e). Unlike the previous comparison of cellular stiffness, the elastic modulus increased substantially with increasing adhesion. Spheroids with *J* = −40 exhibited an elastic modulus of 0.28, increasing to 0.42 for *J* = −50 and 0.52 for *J* = −60. Together, these results indicate that cell-cell adhesion had a substantially greater influence on the elastic modulus of spheroids than the mechanical stiffness of the individual cells from which they were composed.

### Interplay of single-cell stiffness and cell-cell adhesion on the retardation time of spheroids

Both previous spheroid comparisons showed that the fitted retardation time increased despite producing markedly different changes in the elastic modulus. This raises the question of whether cellular stiffness and cell-cell adhesion influence the relaxation dynamics of spheroids independently or jointly. To investigate this, we extended the parameter space to include five values of cell stiffness (*λ_S_* = 0.8, 0.9, 1.0, 1.1, and 1.2) and cell-cell adhesion (*J* = −40, −45, −50, −55, and −60), resulting in 25 distinct spheroid types. For each condition, three forces were simulated (*λ_D_* = 0.6, 0.8, 1.0), and the mean retardation time was summarized as a heat map.

Across all three forces, increasing either the cellular stiffness or the cell-cell adhesion resulted in larger retardation times (Fig. 6). At the lowest force (*λ_D_* = 0.6), *τ* increased monotonically along both parameter axes, reaching its largest values for spheroids composed of the stiffest cells and strongest adhesion (*λ_S_* = 1.2, *J* = −60). The same overall trend was observed for *λ_D_* = 0.8 and 1.0, although the smaller range of retardation times under these larger forces resulted in greater apparent variability. Specifically, *τ* ranged from approximately 7.7–92.8, s for *λ_D_* = 0.6, compared with 3.8–14.1, s and 0.9–7.6, s for *λ_D_* = 0.8 and 1.0, respectively. Together, these results indicate that single-cell stiffness and cell-cell adhesion both contribute to slowing the transition from the initial elastic response to the long-time viscous regime, despite having markedly different effects on the elastic modulus of the spheroid.

**Figure 6:**
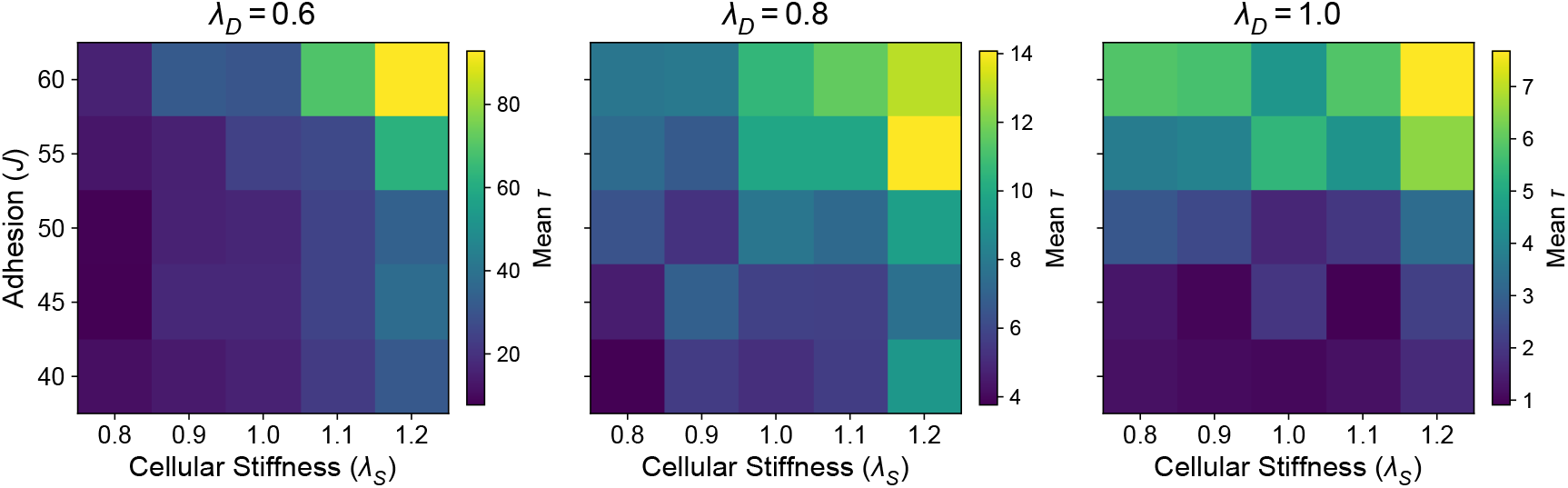
Heat maps of fitted mean retardation time. *τ* . Mean retardation time *τ* as a function of cellular stiffness *λ_S_* and cell-cell adhesion *J* for three forces *λ_D_* = 0.6, 0.8, 1.0.

## Discussion

In this study, we developed a CPM framework for quantitative MPA of both isolated cells and multicellular spheroids within a common computational setting. Using this framework, we investigated how single-cell mechanical properties and cell-cell adhesion contribute to the collective mechanics of spheroids. By independently varying cellular stiffness and cell-cell adhesion, we found that adhesion exerts a substantially stronger influence on the effective elastic modulus of spheroids than the stiffness of the constituent cells themselves. In contrast, both cellular stiffness and cell-cell adhesion contributed to increasing the retardation time governing the transition between the elastic and viscous regimes. Together, these findings suggest that collective mechanical behavior cannot be inferred directly from isolated cell mechanics, but instead depends strongly on intercellular coupling and tissue-level organization.

A central aspect of this work is the representation of MPA within the CPM. Rather than explicitly defining aspiration as an external mechanical force, we represented it through a localized directional motility bias acting only within the pipette region. Although this term formally corresponds cell motility rather than external force application, increasing its magnitude produced progressively larger aspiration and reproduced the characteristic viscoelastic deformation observed experimentally during MPA (Fig. 3c). This phenomenological interpretation allowed the same aspiration protocol to be applied consistently to both isolated cells and spheroids while maintaining a common mechanical framework. More generally, the present framework demonstrates that quantitative aspiration measurements and viscoelastic parameter estimation can be incorporated into the CPM without modifying its underlying stochastic dynamics.

To enable consistent comparisons across simulations performed at different lattice resolutions, we additionally introduced a normalization of the Hamiltonian and *t*_MCS_ with respect to cell size (Fig. S1). Although primarily motivated by the differing computational requirements of single-cell and spheroid simulations, this normalization also facilitates applying the framework to different cell geometries without introducing resolution-dependent mechanical artifacts.

At the single-cell level, increasing the shape penalty strength *λ_S_* consistently reduced aspiration and increased the elastic modulus extracted from aspiration measurements. Likewise, the fitted spring constant *k* increased systematically with *λ_S_*, supporting its interpretation as an effective measure of resistance to deformation within the model. However, *k* also depended on the force, particularly at lower forces, indicating that it should not be interpreted as an intrinsic material constant. Rather, it is interpreted as an effective property where comparisons are most valuable between the same force values. This dependence likely arises from the quadratic form of the surface constraint in Eq. (1), under which energetic penalties increase nonlinearly with increasing deformation. In contrast, the retardation time *τ* decreased with increasing *λ_S_*, indicating faster equilibration in mechanically stiffer cells. Notably, this viscoelastic behavior emerged without explicitly introducing constitutive material equations into the CPM itself. Instead, the combination of stochastic energy minimization together with geometrical area and surface constraints generated aspiration dynamics that were well described by a Kelvin-Voigt model.

Extending the framework from isolated cells to spheroids required additional phenomenological assumptions. Preliminary simulations frequently exhibited either prolonged stochastic waiting periods before aspiration or immediate, nearly unrestricted aspiration following entry into the pipette. To obtain stable aspiration dynamics over experimentally relevant timescales, peripheral cells were assigned a reduced shape penalty to facilitate aspiration initiation while preserving that of the spheroid interior intact. In addition, Bell-like catch-bond dynamics were introduced by allowing cell-cell adhesion to increase locally within the pipette. Together, these modifications reproduced the characteristic transition from an initial elastic deformation to prolonged viscous creep described by a three-element viscoelastic model. Although these assumptions remain phenomenological and should not be interpreted as explicit molecular mechanisms, they provided a reproducible computational framework for systematically comparing the effects of cellular stiffness and cell-cell adhesion on multicellular mechanics.

Perhaps the most notable result of this study is that increasing cell stiffness had little influence on the elastic modulus of spheroids despite substantially altering the mechanics of isolated cells. Although spheroids composed of stiffer cells exhibited a higher force threshold for initial aspiration, their elastic modulus remained nearly unchanged across the investigated *λ_S_* range. The fitted spring constant *k* showed similarly modest variation between the different spheroid conditions, consistent with the elastic modulus. In contrast, varying cell-cell adhesion produced pronounced changes in both aspiration dynamics and the elastic modulus. Within the present framework, these findings suggest that the elastic response of spheroids is governed primarily by the mechanical coupling between neighboring cells rather than by the mechanical stiffness of the constituent cells alone.

The retardation time exhibited a different dependence on model parameters. Unlike the elastic modulus, which was primarily influenced by cell-cell adhesion, the retardation time increased with both cellular stiffness and cell-cell adhesion. This indicates that these two mechanical descriptors capture distinct aspects of spheroid mechanics. Whereas the elastic modulus characterizes the magnitude of the elastic deformation, the retardation time reflects the timescale over which the spheroid transitions from the initial elastic response toward long-time viscous flow. The persistence of this trend across a broader parameter space suggests that cellular stiffness and cell-cell adhesion jointly influence collective relaxation dynamics, despite contributing differently to the elastic response itself.

The opposite dependence of the retardation time at the single-cell and spheroid scales likely reflects differences in the processes represented by the fitted transient. For an isolated cell, increasing the shape penalty strength increased *k* while reducing *τ*, the time required to approach the aspiration plateau. In the Kelvin-Voigt representation, *τ* = *ξ_f_ /k*, suggesting that the increase in *k* exceeded any corresponding change in effective dissipation. On the other hand, spheroid aspiration requires additional processes, such as cell jamming and load transmission through adhesive contacts. Increasing cellular stiffness may hinder these collective configurational changes, thereby increasing *ξ_f_* sufficiently to produce a longer *τ*, even when *k* changes little.

Finally, our model has several limitations. Although isolated cells and spheroids were examined within a common CPM framework, the mechanical parameters extracted at the two scales should not be interpreted as direct measurements of the same constitutive material property. Spheroid aspiration additionally reflects cell-cell adhesion, collective rearrangement, and load redistribution, whereas single-cell aspiration primarily reflects deformation from a target shape. Moreover, the effective force parameter *λ_D_* was not calibrated to a physical pressure or force. Consequently, the fitted spring constants and effective elastic moduli were reported in CPM units and should be interpreted as relative mechanical indices within the simulation framework rather than absolute material properties. The spheroid model also includes phenomenological assumptions, including peripheral softening and local adhesion strengthening within the pipette, which facilitate experimentally consistent aspiration dynamics but do not represent uniquely identified molecular mechanisms. Additionally, the simulations were performed in two dimensions and therefore do not capture the full three-dimensional geometry of experimental spheroids. Despite these limitations, the model provides a unified and experimentally informed framework for comparing how cellular mechanics and intercellular adhesion shape multicellular viscoelastic behavior across scales.

## Author Contributions

D.D. and P.A.I. designed the research. P.A.I. and D.N.R. supervised the research. D.D. performed the simulations and analyzed the results. E.P. performed the experiments. D.D. analyzed microscopy images. D.D. wrote the draft. All authors reviewed and edited the manuscript.

## Acknowledgments

Acknowledge assistance that does not meet the criteria for authorship. Confirm that acknowledged individuals consent to being named where required.

## Supplementary Information

### Cell size scaling

#### Scaling of the Hamiltonian terms

In the Cellular Potts Model, copy acceptance is governed by changes in the Hamiltonian. A force-like quantity may be defined by differentiating the Hamiltonian with respect to a configurational coordinate **q**,

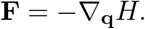

The derivatives considered below are instead sensitivities to the target geometric parameters and should not be interpreted directly as mechanical forces.

Consider first the perimeter contribution for a given cell,

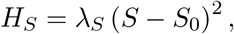

where *S* is the cell perimeter,

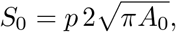

*A*_0_ is the target area, and *p* is a dimensionless shape parameter. The case *p* = 1 corresponds to a circle.

The sensitivity to the target perimeter is

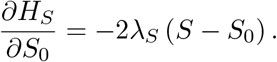

Assume geometrically similar cells with fixed *p* and a fixed relative perimeter mismatch as the cell size changes. Since

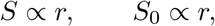

it follows that

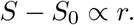

Consequently,

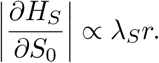

Let *r*_ref_ denote the reference radius at which *λ_S_* was calibrated, and let *r*_current_ denote the radius of the rescaled cell. Preserving the magnitude of this sensitivity requires

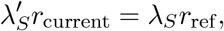

and therefore

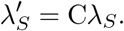

where C = *r*_ref_ */r*_current_.

The area contribution is

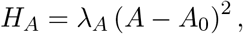

with sensitivity

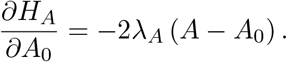

For geometrically similar cells with a fixed relative area mismatch,

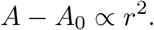

Hence,

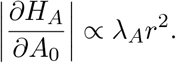

Preserving this sensitivity gives

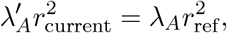

so that

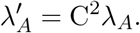

The directed-motion contribution is

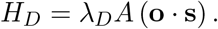

Assuming that the directional factor is dimensionless and remains of order unity, the total contribution scales as

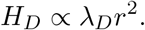

Therefore, preserving the magnitude of the total directed-motion contribution requires

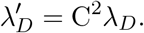

#### Monte Carlo time scaling

We performed a sensitivity analysis to determine whether the *t*_MCS_ depends on the total number of lattice sites or on the number of occupied pixels (Fig. S2). Increasing the grid size from *N* × *N* to 2*N* × 2*N*, while keeping the cell and all model parameters unchanged, produced nearly identical process timescales. Scaling *t*_MCS_ duration by the fourfold increase in total lattice area instead made the dynamics substantially slower.

These results indicate that the relevant scaling is determined by the number of occupied pixels rather than by the total number of lattice sites. In the present case, where the occupied region consists of a single two-dimensional cell,

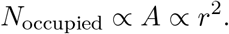

The scaled simulation time is therefore

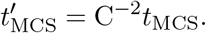

**Figure S1:**
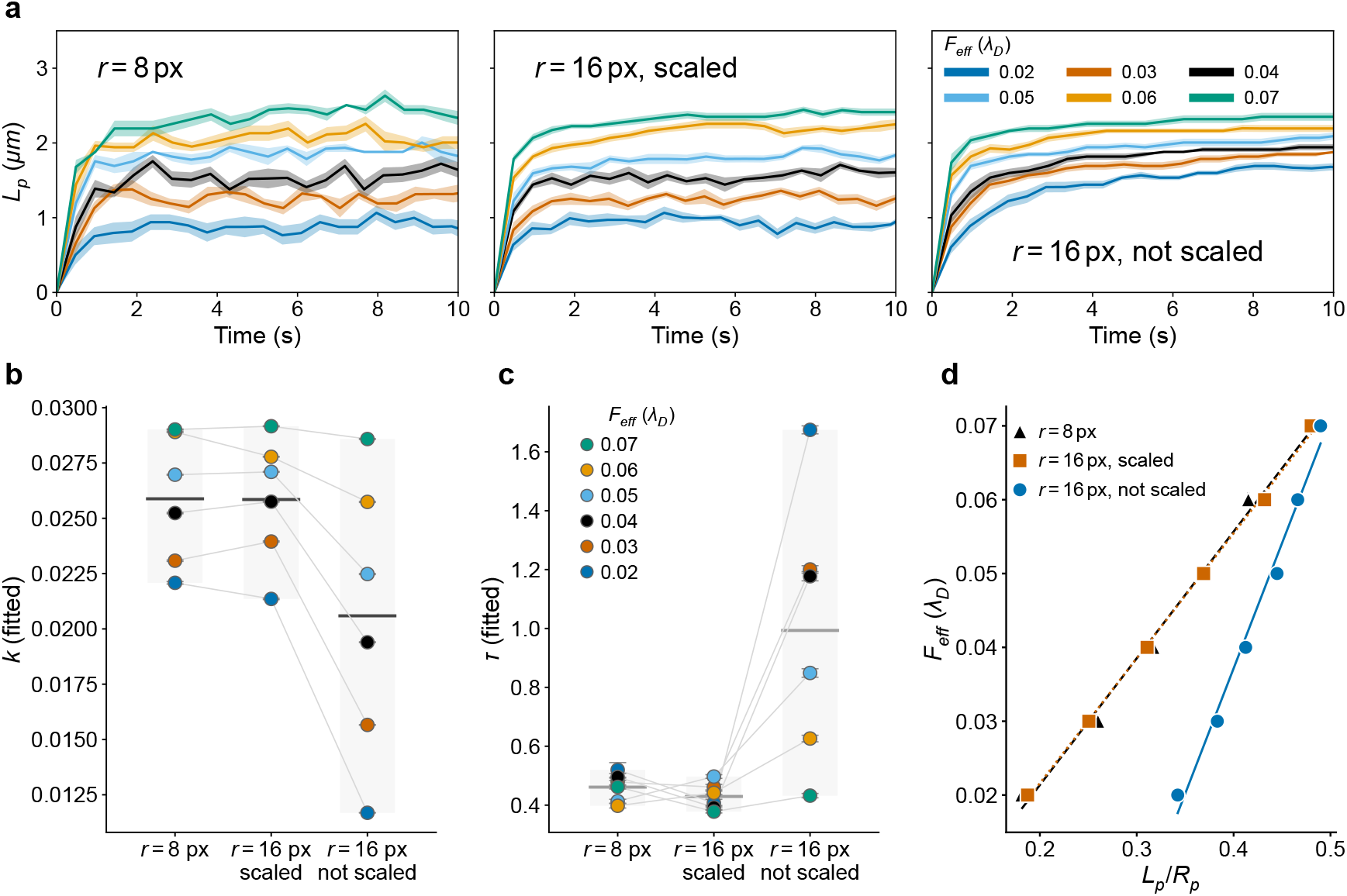
Single-cell mechanics for scaled and unscaled cells using the scaling convention. **a.** Single-cell aspiration length *L_p_* over time for six values of the effective aspiration force parameter *λ_D_* for a reference cell with *r* = 2.5 *µ*m (left), a cell with *r* = 5 *µ*m scaled to the reference cell (middle), and a cell with *r* = 5 *µ*m with no scaling (right). Grid sizes were proportional to the cell size in all three cases, resulting in effective resolution change alone. **b**. Fitted spring parameter *k* for the three conditions. Gray lines connect values obtained at the same effective force. **c**. Fitted retardation time *τ* for the three conditions. Gray lines connect values obtained at the same effective force. **d**. Effective aspiration force as a function of normalized aspiration length *L_p_/R_p_* for the three conditions. Linear regression slopes are proportional to the effective elastic modulus, 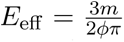. Effective elastic moduli are 0.039 for the cell with *r* = 2.5 *µ*m, 0.038 for the scaled cell with *r* = 5 *µ*m, and 0.077 for the unscaled cell with *r* = 5 *µ*m. Error bars indicate 95% confidence intervals estimated from nonlinear least-squares regression.

**Figure S2:**
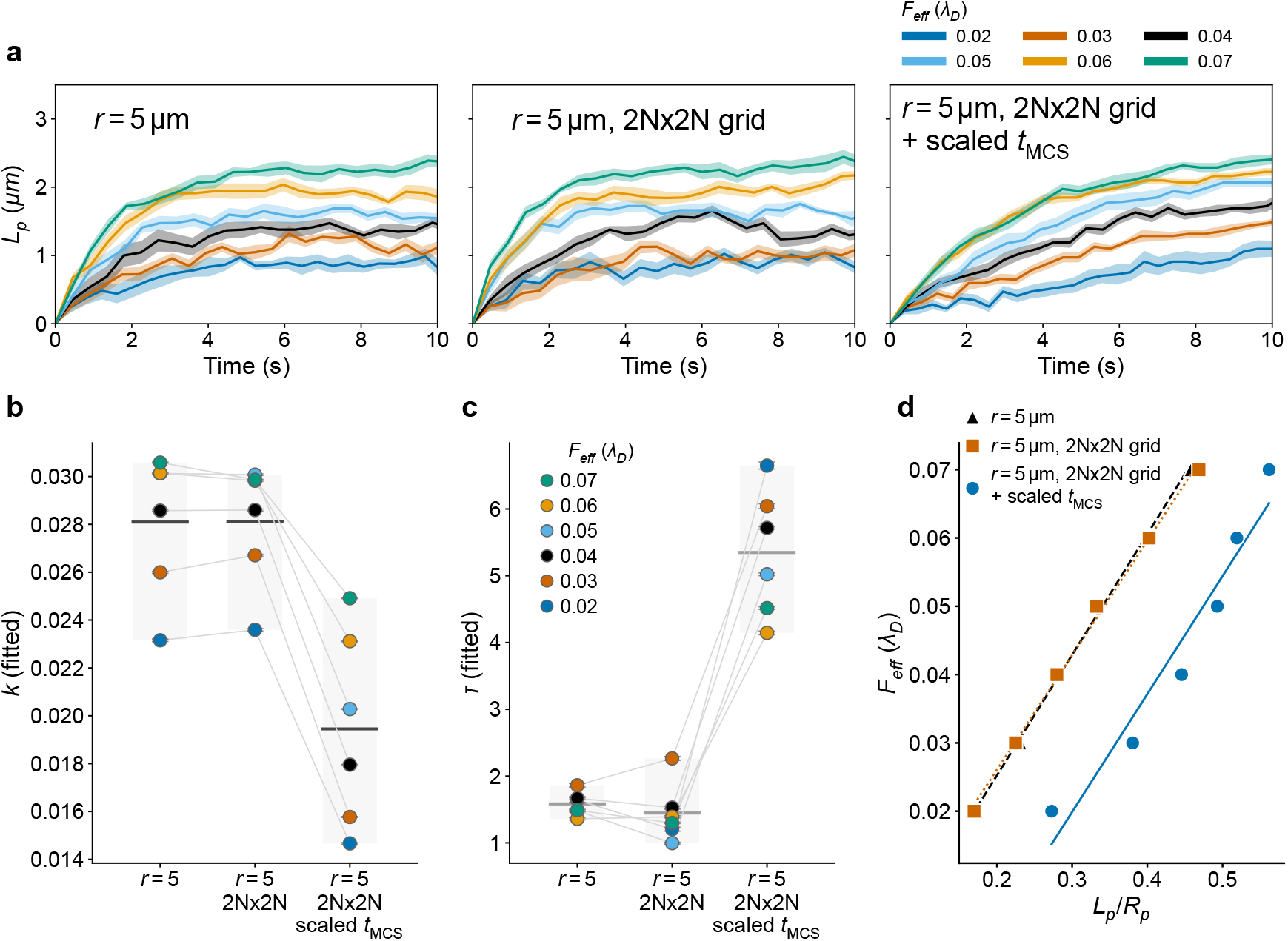
Sensitivity analysis to lattice size. **a**. Single-cell aspiration length *L_p_* over time for six values of the effective aspiration force parameter *λ_D_*for a reference cell with *r* = 5 *µ*m (left), a cell with *r* = 5 *µ*m inside a lattice of double area (middle), and a cell with *r* = 5 *µ*m inside a lattice of double area with scaled MCS time (right). All three cells had a radius of *r* = 16 px and were scaled to a reference cell with *r* = 5 px using the Hamiltonian scaling. **b**. Fitted spring parameter *k* for the three conditions. Gray lines connect values obtained at the same effective force. **c**. Fitted retardation time *τ* for the three conditions. Gray lines connect values obtained at the same effective force. **d**. Effective aspiration force as a function of normalized aspiration length *L_p_/R_p_* for the three conditions. Linear regression slopes are proportional to the effective elastic modulus, 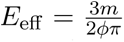. Effective elastic moduli are 0.04 for the cell with *r* = 2.5 *µ*m, 0.038 for the scaled cell with *r* = 5 *µ*m, and 0.039 for the unscaled cell with *r* = 5 *µ*m. Error bars indicate 95% confidence intervals estimated from nonlinear least-squares regression.

**Figure S3:**
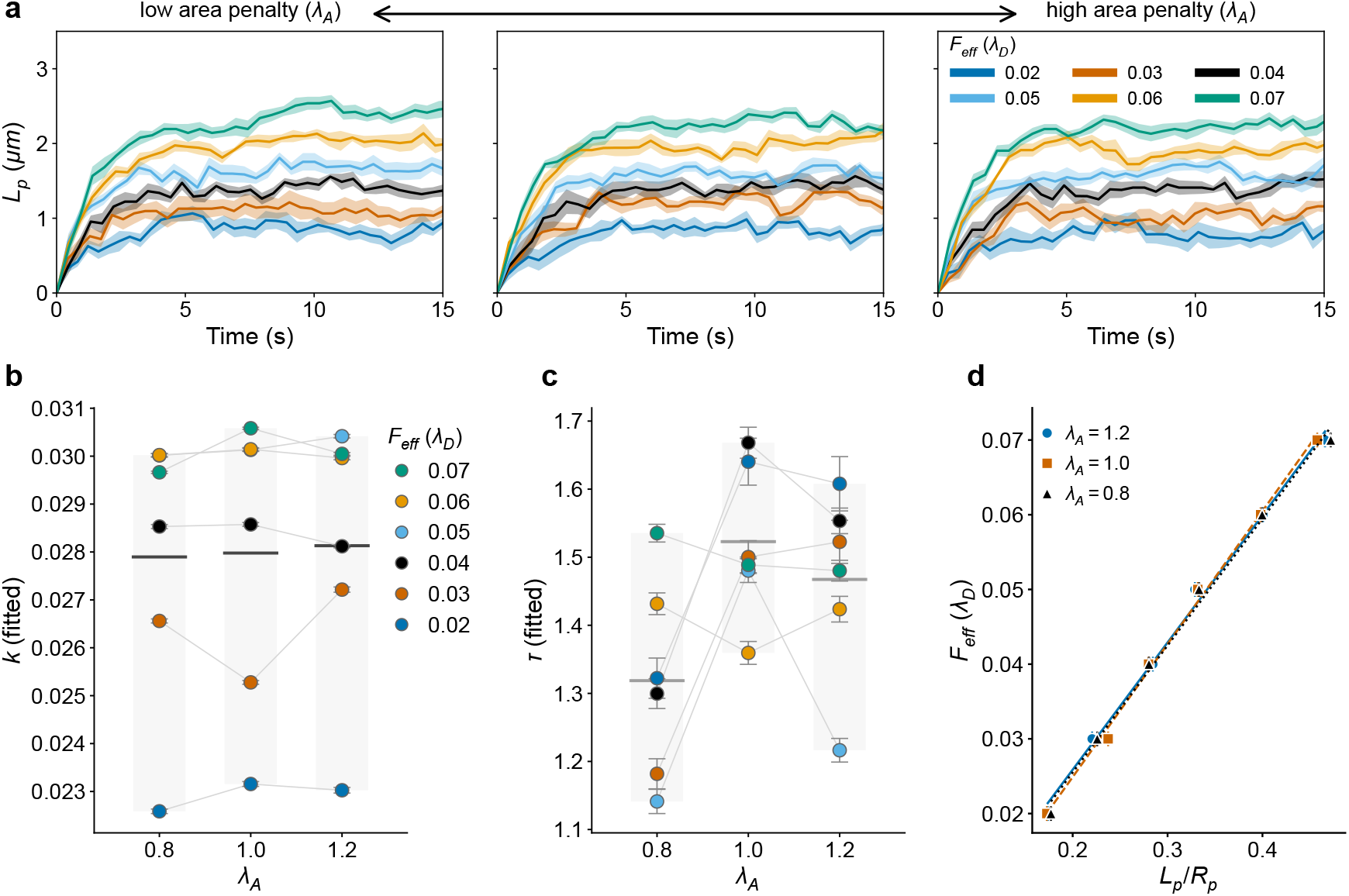
Single-cell mechanics controlled by the area constraint strength. *λ_A_***. a**. Single-cell aspiration length *L_p_* over time for six values of the effective aspiration force parameter *λ_D_* across three values of area constraint strength *λ_A_*: 0.8 (left), 1.0 (middle), and 1.2 (right). **b**. Fitted spring parameter *k* for the three *λ_A_* values. Gray lines connect values obtained at the same effective force. **c**. Fitted retardation time *τ* for the three *λ_A_* values. Gray lines connect values obtained at the same effective force. **d**. Effective aspiration force as a function of normalized aspiration length *L_p_/R_p_* for the three *λ_A_* values. Linear regression slopes are proportional to the effective elastic modulus, 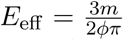 . Effective elastic moduli are 0.039 for both *λ_A_*= 0.8 and *λ_A_*= 1.2, and 0.04 for *λ_A_* = 1.0. Error bars indicate 95% confidence intervals estimated from nonlinear least-squares regression.

**Table S1:** Simulation parameters.

| Parameter | Value | Description |
| --- | --- | --- |
| Lattice size (cell) | (161 px, 260 px, 0), 1 px $\approx 0.3 \mu\text{m}$ | $(x, y, z)$ coordinates; 10 cells aspirated |
| Lattice size (spheroid) | (410 px, 250 px, 0), 1 px $\approx 1 \mu\text{m}$ | $(x, y, z)$ coordinates |
| $T$ | 0.2 | Temperature |
| $t_{\text{MCS}}$ | 0.1 | Model time corresponding to one MCS |
| $\lambda_A$ | 1.0 | Strength of the area constraint |
| $A_0$ | 80.0 px | Target cell area |
| $\lambda_S$ | [0.8, 0.9, 1.0, 1.1, 1.2] | Strength of the shape constraint |
| $\lambda_D$ (cell) | [0.02, 0.03, 0.04, 0.05, 0.06, 0.07] | Effective aspiration force strength |
| $\lambda_D$ (spheroid) | [0.6, 0.7, 0.8, 0.9, 1.0] | Effective aspiration force strength |
| $p$ | 1.0 | Target asphericity |
| $J_{\text{CM}}$ | 0.0 | Cell-medium adhesion |
| $J_{\text{CC}}$ | [−40, −45, −50, −55, −60] | Cell-cell adhesion |
| $N_C$ | 500 | Number of cells in one spheroid |
| $R_p$ (cell) | 8.0 px ($5.0 \mu\text{m}$) | Pipette radius |
| $R_p$ (spheroid) | 100.0 px ($15.0 \mu\text{m}$) | Pipette radius |

